# Sleep prevents catastrophic forgetting in spiking neural networks by forming joint synaptic weight representations

**DOI:** 10.1101/688622

**Authors:** Ryan Golden, Jean Erik Delanois, Pavel Sanda, Maxim Bazhenov

## Abstract

Artificial neural networks overwrite previously learned tasks when trained sequentially, a phenomenon known as catastrophic forgetting. In contrast, the brain learns continuously, and typically learns best when new learning is interleaved with periods of sleep for memory consolidation. In this study, we used spiking network to study mechanisms behind catastrophic forgetting and the role of sleep in preventing it. The network could be trained to learn a complex foraging task but exhibited catastrophic forgetting when trained sequentially on multiple tasks. New task training moved the synaptic weight configuration away from the manifold representing old tasks leading to forgetting. Interleaving new task training with periods of off-line reactivation, mimicking biological sleep, mitigated catastrophic forgetting by pushing the synaptic weight configuration towards the intersection of the solution manifolds representing multiple tasks. The study reveals a possible strategy of synaptic weights dynamics the brain applies during sleep to prevent forgetting and optimize learning.

## Introduction

Humans are capable of continuously learning to perform novel tasks throughout life without interfering with their ability to perform previous tasks. Conversely, while modern artificial neural networks (ANNs) are capable of learning to perform complicated tasks, ANNs have difficulty learning multiple tasks sequentially^1–3^. Sequential training commonly results in catastrophic forgetting, a phenomenon which occurs when training on the new task completely overwrites the synaptic weights learned during the previous task, leaving the ANN incapable of performing a previous task^1–4^. Attempts to solve catastrophic forgetting have drawn on insights from the study of neurobiological learning, leading to the growth of neuroscience-inspired artificial intelligence (AI)^5–7^. While these approaches are capable of mitigating catastrophic forgetting in certain circumstances^6^, a general solution which can achieve human level performance for continual learning is still an open question.

Historically, an interleaved training paradigm, where multiple tasks are presented within a common training dataset, has been employed to circumvent the issue of catastrophic forgetting^4,8,9^. In fact, interleaved training was originally construed to be an approximation to what the brain may be doing during sleep to consolidate memories; spontaneously reactivating memories from multiple interfering tasks in an interleaved manner^9^. Unfortunately, explicit use of interleaved training, in contrast to memory consolidation during biological sleep, imposes the stringent constraint that the original training data be perpetually stored for later use and combined with new data to retrain the network^1,2,4,9^. Thus, the challenge is to understand how the biological brain enables memory reactivation during sleep without access to past training data.

Parallel to the growth of neuroscience-inspired ANNs, there has been increasing investigation of spiking neural networks (SNNs) which attempt to provide a more realistic model of brain functioning by taking into account the underlying neural dynamics and by using biologically plausible local learning rules^10–13^. A potential advantage of the SNNs, that was explored in our new study, is that local learning rules combined with spike-based communication allow previously learned memory traces to reactivate spontaneously and without interference during off-line processing – sleep. A common hypothesis, supported by a vast range of neuroscience data, is that the consolidation of memories during sleep occurs through local unsupervised synaptic changes enabled by reactivation of the neuron ensembles engaged during learning^14^. Indeed, spike sequence replay was observed in the neocortex^15–17^ following both hippocampal-dependent tasks^15^ and hippocampal-independent tasks^18^.

Here we used a multi-layer SNN with reinforcement learning to investigate whether interleaving periods of new task training with periods of noise-induced spontaneous reactivation, resembling sleep in the brain^19–21^, can circumvent catastrophic forgetting. The network could be trained to learn one of two complementary complex foraging tasks involving pattern discrimination but exhibits catastrophic forgetting when trained on the tasks sequentially. Significantly, we show that catastrophic forgetting can be prevented by periodically interrupting reinforcement learning on a new task with unsupervised sleep phases. While new task training alone moved synaptic weight configuration away from the solution manifold representing old tasks and towards the manifold specific for new task, interleaving new task training with unsupervised sleep replay allowed the synaptic weights to stay near the manifold specific for the old task and still to move towards its intersection with the manifold representing the new task. Our study predicts that sleep prevents catastrophic forgetting in the brain by forming joint synaptic weight representations suitable for storing multiple memories.

## Results

### Complementary complex foraging tasks can be robustly learned

We modeled a simple 3-layer feedforward spiking neural network (see Figure 1A and *Methods: Network Structure* for details) simulating basic steps from sensory input to motor output in the brain. Excitatory synapses between the input (I) and hidden (H) layers were subjected to unsupervised learning (implemented as non-rewarded STDP)^22,23^ while those between the H and output (O) layers were subjected to reinforcement learning (implemented using rewarded STDP)^24–27^ (see *Methods: Synaptic plasticity* for details). Unsupervised plasticity allowed neurons in layer H to learn different particle patterns at various spatial locations of the input layer I, while rewarded STDP allowed the neurons in layer O to learn motor decisions based on the type of the particle patterns detected in the visual field^12^. We trained the network on one of two complementary complex foraging tasks. In either task, the network learned to discriminate between a rewarded and a punished particle pattern in order to acquire as much of the rewarded patterns as possible. In the following we consider pattern discriminability (rewarded vs punished) as a measure of performance, with chance performance being 0.5.

**Figure 1.**
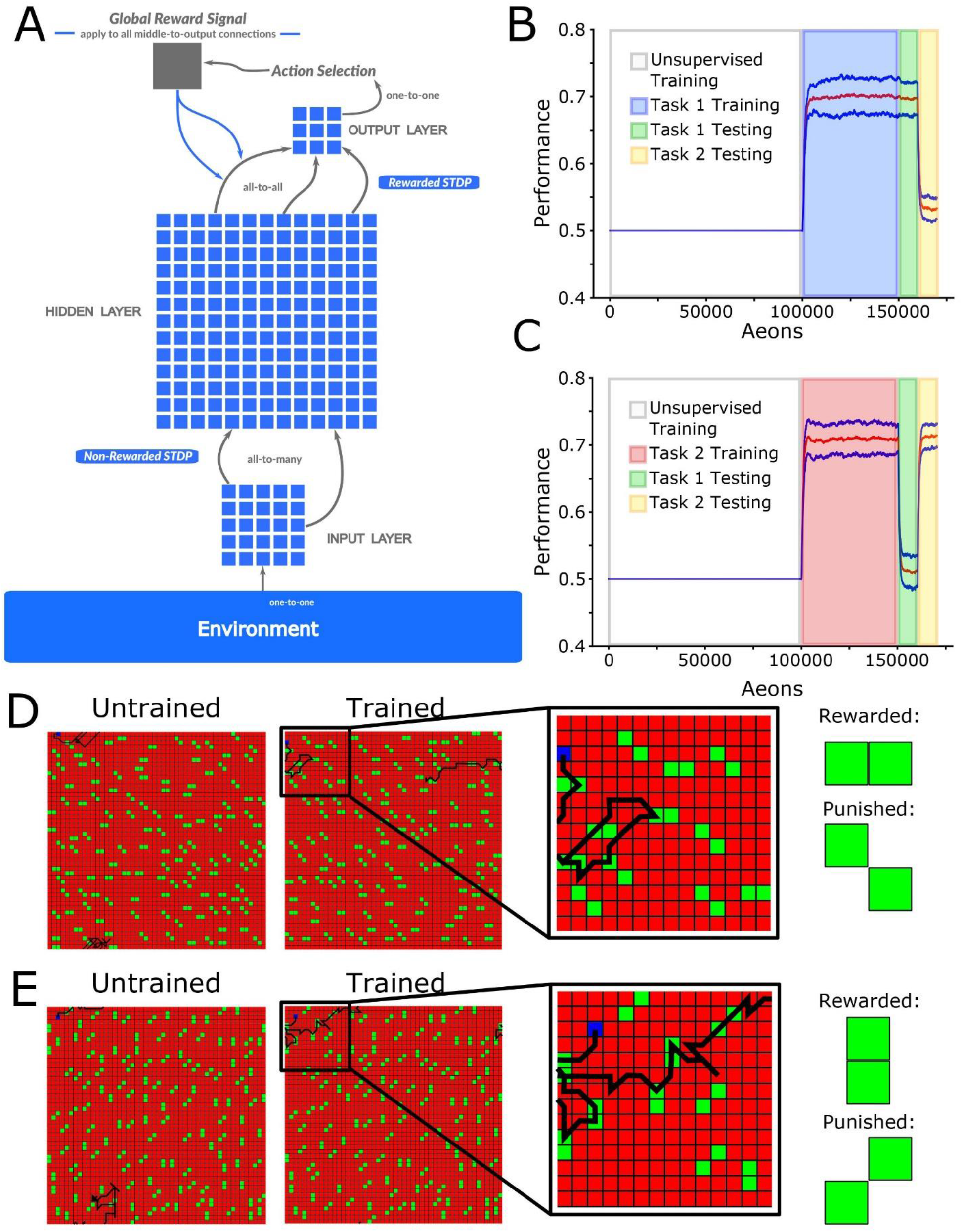
Network architecture and complementary foraging task structure. **(A)** The network had three layers of neurons with a feed-forward connectivity scheme. Input from the “visual field” (7×7 subspace of 50×50 virtual environment) was simulated as a set of excitatory inputs to the input layer neurons representing the position of food particles in an egocentric reference frame relative to the virtual agent. Each hidden layer neuron received an excitatory synapse from 9 randomly selected input layer neurons. Excitatory synapses between input and hidden layer neurons were subject to unsupervised STDP, while those between hidden and output layer neurons were subject to rewarded STDP. Each output layer neuron received one excitatory and one inhibitory synapse from each hidden layer neuron. The most active neuron in the output layer (size 3×3) determined the direction of movement. **(B)** Mean performance (red line) and standard deviation (blue lines) over time: 100,000 aeons (1 aeon = 100 movement cycles) of unsupervised training (white), 50,000 aeons of Task 1 training (blue), and 10,000 aeons of Task 1 (green) and Task 2 (yellow) testing. The y-axis represents the agent’s performance, or the probability of acquiring rewarded as opposed to punished particle patterns. The x-axis is time in aeons. Mean performance during testing on Task 1 was 0.70 ± 0.02 while Task 2 was 0.53 ± 0.02. **(C)** The same as shown in (B) except now for: 10,000 aeons of unsupervised training (white), 5000 aeons of Task 2 training (red), and 1,000 aeons of Task 1 (green) and Task 2 (yellow) testing. Mean performance during testing on Task 1 was 0.51 ± 0.02 while Task 2 was 0.71 ± 0.02. **(D)** Examples of trajectories through the environment at the beginning (left) and at the end (middle-left) of training on Task 1, with a zoom in on the trajectory at the end of training (middle-right), and the values of the task-relevant food particles (right). **(E)**. The same as shown in (D) except now for Task 2.

The paradigm for Task 1 is shown in Figure 1B. First, during an unsupervised learning period, all 4 types of 2-particle patterns (horizontal, vertical, positive diagonal, and negative diagonal) were present in the environment with equal densities. This was a period, equivalent to a developmental critical period in the brain, when the network learned the environmental statistics and formed, in layer H, high level representation of all possible patterns found at the different visual field locations (see Figure 2 for details). Unsupervised training was followed by a reinforcement learning period, equivalent to task specific training in the brain, during which the synapses between layers I and H were frozen but synapses from H to O were updated using a rewarded STDP rule. The reinforcement learning period was when the network learned to make decisions about which direction to move based on the visual input. Whether patterns were rewarded during reinforcement learning depended on the task - for Task 1 horizontal patterns were rewarded and negative diagonal patterns were punished (Figure 1D). During both the rewarded training and the testing periods only 2 types of patterns were present in the environment (e.g. horizontal and negative diagonal for Task 1).

**Figure 2.**
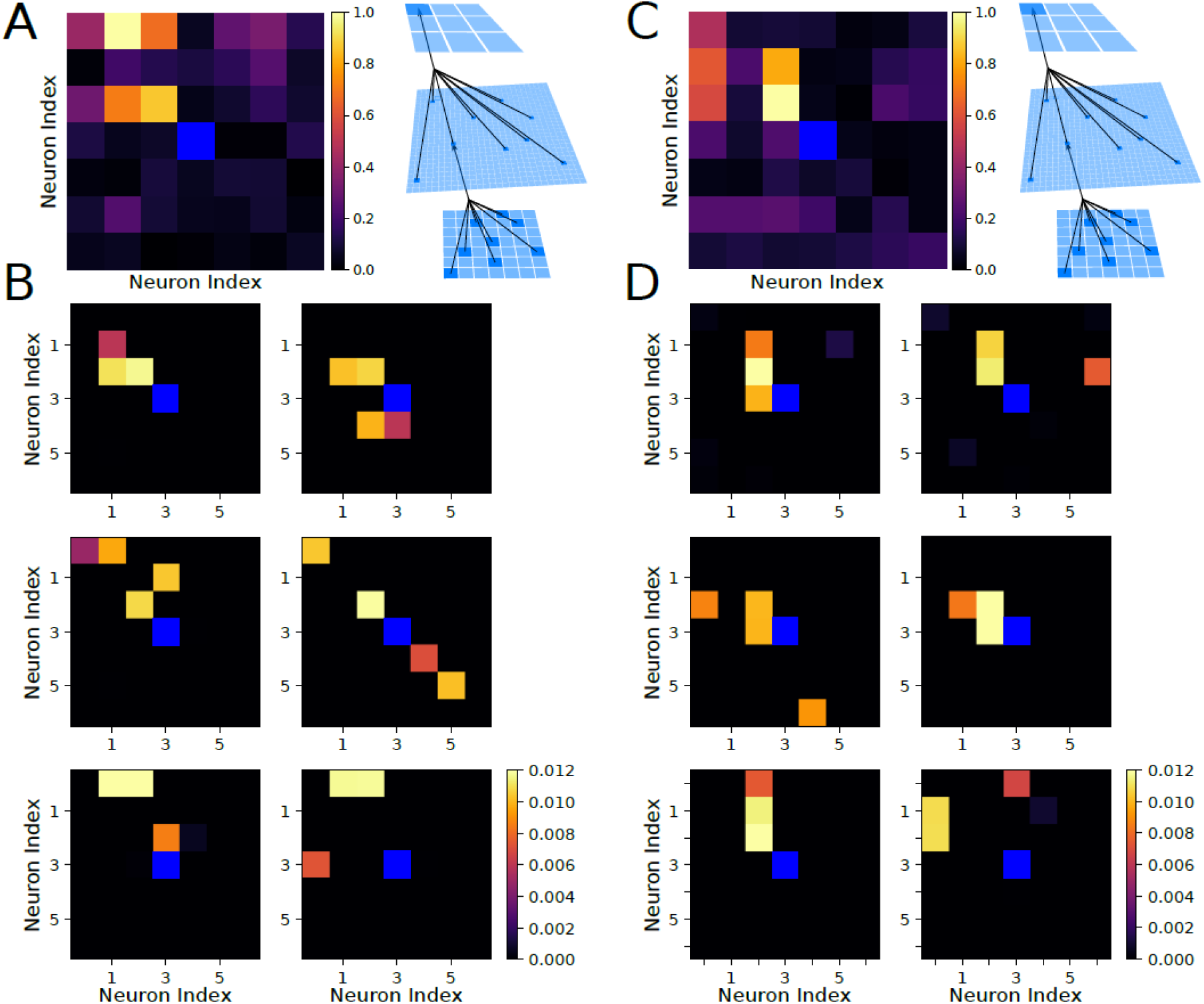
Receptive fields of output and hidden layer neurons determine the agent policy. **(A)** Left, Receptive field of the output layer neuron controlling movement to the upper-left direction following training on Task 1. This neuron can be seen to selectively respond to horizontal orientations in the upper-left quadrant of the visual field. Right, Schematic of connections between layers. **(B)** Examples of receptive fields of hidden layer neurons which synapse strongly onto the output neuron from (A) after training on Task 1. The majority of these neurons selectively respond to horizontal food particles in the upper-left quadrant of the visual field, with one neuron (middle-right) selectively responding to the presence of negative diagonal food particles in the bottom-right quadrant and the lack of negative diagonal food particles in the upper-left quadrant of the visual field. **(C)** The same as shown in (A) except following training on Task 2. The upper-left decision neuron can be seen to selectively respond to vertical orientations in the upper-left quadrant of the visual field. **(D)** The same as shown in (B) except following training on Task 2. All of these neurons selectively respond to vertical food particles in the upper-left quadrant of the visual field.

After training Task 1, mean performance on Task 1 was 0.70 ± 0.2 while on Task 2 (which has not been trained yet) was 0.53 ± 0.2 (chance level). Figure 1D shows examples of trajectories of the simulated agent at the beginning of (left) and after (right) reinforcement learning period. The naive agent moved randomly through the environment, but after training it moved to seek out horizontal patterns and largely avoid negative diagonal ones. The complementary paradigm for Task 2 (vertical patterns are rewarded and positive diagonal are punished) is shown in Figure 1C,E. These results demonstrate that the network is capable of learning and performing either one of the two complementary complex foraging tasks.

To get an understanding of the policy developed by the network for each task, we computed the receptive field of each neuron in layer O with respect to the input from layer I (see schematic in Figures 2A/C). This was done by first computing the receptive fields of all of the neurons in layer H with respect to I, then performing a weighted average where the weights were given by the synaptic strength from each neuron in layer H to the particular neuron in layer O. Figure 2A shows a representative example of the receptive field which developed after training on Task 1 for one specific neuron in layer O which controls movements to the upper-left direction. This neuron responded most robustly to bars of horizontal orientation (rewarded) in the upper-left quadrant of the visual field and, importantly, did not respond to bars of negative diagonal orientation (punished).

Figure 2B shows examples of receptive fields of six neurons in layer H which synapse strongly onto the upper-left neuron in layer O (the neuron shown in Figure 2A). These neurons form high level representations of the input patterns, similar to the neurons in the higher levels of the visual system or later layers of a convolutional neural network^28–30^. The majority of these receptive fields revealed strong selection for the horizontal (i.e. rewarded) food particles in the upper-left quadrant of the visual field. As a particularly notable example, one of these layer H neurons (Figure 2B; middle-right) preferentially responded to negative diagonal (i.e. punished) food particles in the bottom-right quadrant of the visual field. Thus, spiking in this neuron caused the agent to move away from these punished food particles. Similar findings after training on Task 2 are shown in Figures 2C and 2D.

### Catastrophic forgetting occurs following sequential but not interleaved training

We next tested whether the network model could exhibit catastrophic forgetting by training sequentially on Task 1 (old task here) followed by Task 2 (new task) (Figure 3A). Following Task 2 training, performance on Task 1 was down to no better than chance (0.52 ± 0.02), while performance on Task 2 improved to 0.69 ± 0.03 (Figure 3 A,B). Thus, sequential training on a complementary task caused the network to undergo catastrophic forgetting of the task trained earlier, remembering only the most recent task.

**Figure 3.**
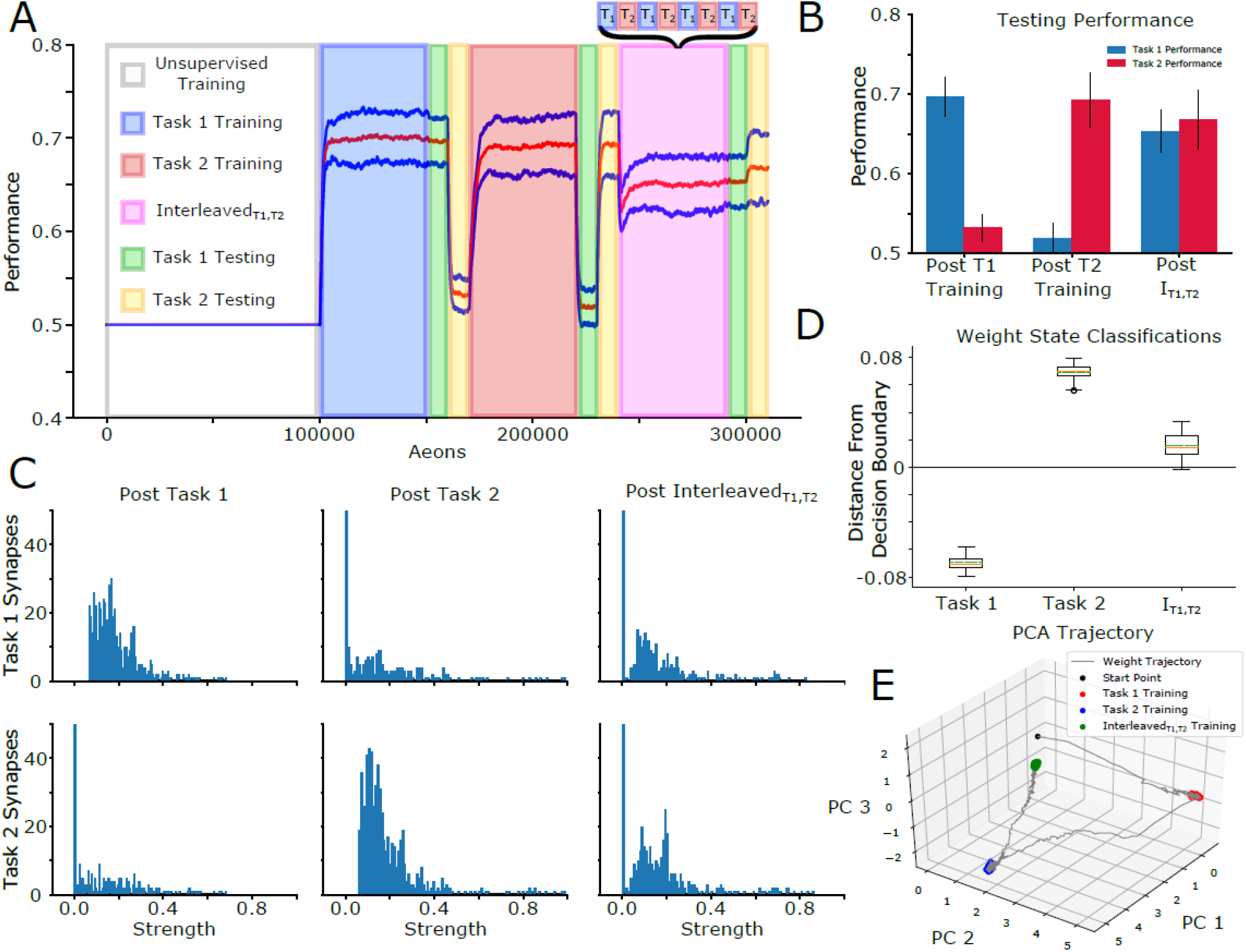
Sequential training on complementary tasks induces catastrophic forgetting which can be rescued by interleaved training. **(A)** Mean performance (red line) and standard deviation (blue lines) over time: 100,000 aeons of unsupervised training (white), 50,000 aeons of Task 1 training (blue), 10,000 aeons of Task 1 (green) and Task 2 (yellow) testing, 50,000 aeons of Task 2 training (red), 10,000 aeons of Task 1 (green) and Task 2 (yellow) testing, 50,000 aeons of Interleaved_T1,T2_ training (purple), 10,000 aeons of Task 1 (green) and Task 2 (yellow) testing. **(B)** Mean and standard deviation of performance during testing on Task 1 (blue) and Task 2 (red) after each training period. Following Task 1 training, mean performance on Task 1 was 0.69 ± 0.02 while Task 2 was 0.53 ± 0.02. Conversely, following Task 2 training, mean performance on Task 1 was 0.52 ± 0.02 while Task 2 was 0.69 ± 0.04. Following Interleaved_T1,T2_ training, mean performance on Task 1 was 0.65 ± 0.03 while Task 2 was 0.67 ± 0.04. **(C)** Distributions of task-relevant synaptic weights. The distributional structure of Task 1-relevant synapses following Task 1 training (top-left) is destroyed following Task 2 training (top-middle), but partially recovered following Interleaved_T1,T2_ training (top-right). Similarly, the distributional structure of Task 2-relevant synapses following Task 2 training (bottom-middle), which was not present following Task 1 training (bottom-left), was partially preserved following Interleaved_T1,T2_ training (bottom-right). **(D)** Box plots with mean (dashed green line) and median (dashed orange line) of the distance to the decision boundary found by an SVM trained to classify Task 1 and Task 2 synaptic weight matrices for Task 1, Task 2, and Interleaved_T1,T2_ training across trials. Task 1 and Task 2 synaptic weight matrices had mean classification values of −0.069 and 0.069 respectively, while that of Interleaved_T1,T2_ training was 0.016. **(E)** Trajectory of H to O layer synaptic weights through PC space. Synaptic weights which evolved during Interleaved_T1,T2_ training (green dots) clustered in a location of PC space intermediary between the clusters of synaptic weights which evolved during training on Task 1 (red dots) and Task 2 (blue dots).

Interleaved training was proposed as a solution for catastrophic forgetting^4,8,9^, so we added an Interleaved Task 1 and Task 2 (Interleaved_T1,T2_) training phase to our simulation (Figure 3A) to test whether it was a capable of learning Task 1 (now new task) without overwriting Task 2 (old task). For interleaved training we alternated short presentations of Task 1 and Task 2 every 100 movement cycles. Figure 3B shows that, following Interleaved_T1,T2_ training, the network achieved a performance of 0.65 ± 0.03 on Task 1 and a performance of 0.67 ± 0.04 on Task 2. Therefore, Interleaved_T1,T2_ training allowed the network to relearn Task 1 without forgetting what the network had just learned during training on Task 2. Note, we also tested Interleaved_T1,T2_ training right after the unsupervised phase and found the same high performance for both Task 1 and Task 2 (not shown).

We identified task-relevant synapses after training on a given task (top 10% of synapses), and we traced the same set of synapses after training on the opposing task or after Interleavedψ1,T2 training. The structure in the distribution of Task 1-relevant synapses following Task 1 training (Figure 3C, top-left) was destroyed following Task 2 training (top-middle; i.e., majority of Task 1-relevant synapses were reduced to zero after Task 2 training) but partially recovered following Interleaved_T1,T2_ training (top-right). Similarly, the structure in the distribution of Task 2-relevant synapses following Task 2 training (bottom-middle) was not present following Task 1 training (bottom-left) and was partially retained following Interleaved_T1,T2_ training (bottom-right).

To better understand the effect of Interleaved_T1,T2_ training on the synaptic weights, we trained a support vector machine (SVM; see *Method: Support Vector Machine Training* for details) with a radial basis function kernel to classify the synaptic weight configurations between layers H and O (i.e. those responsible for decision making) according to whether they serve to perform Task 1 or Task 2. Figure 3D shows the average distance from the decision boundary across trials for synaptic weights associated with Task 1, Task 2, and Interleaved_T1,T2_ training. While the SVM robustly classified the synaptic weight matrices from Task 1 and Task 2, the weight states after Interleaved_T1,T2_ training were significantly closer to the decision boundary (typically on the task 2 side). This indicates that the synaptic weight matrices from Interleaved_T1,T2_ training are a mixture of Task 1 and Task 2 states.

Figure 3E shows the trajectory of the synaptic weight distribution for the experiment in Figure 3A projected to 3-dimensions using principal components analysis (PCA). It can be seen that while synaptic weight matrices associated with Task 1 and Task 2 training cluster in distinct regions of PC space, Interleaved_T1,T2_ training pushes the synaptic weights to an intermediate location between Task 1 and Task 2.

### Periods of sleep allow for sequential training without catastrophic forgetting

Sleep is believed to be an off-line processing period when recent memories are replayed to avoid damage by new learning. Particularly for procedural (hippocampal-independent) memories, rapid-eye-movement (REM) sleep may organize neuronal activity to replay memory traces^31^. Can we implement a sleep like phase to our model to protect an old task and still accomplish new task learning without explicit re-training of the old task (e.g., without doing explicit interleaved training of Task 1 and Task 2)?

Again, we first trained the network on Task 1 and Task 2 sequentially to illustrate occurrence of catastrophic forgetting (Figure 4A). At this point the network remembered the most recent task (i.e. Task 2) but Task 1 was forgotten. Next, we implemented a training phase consisted of alternating periods of training on Task 1 (considered to be a new task here) lasting 100 movement cycles and periods of “sleep” of the same duration (we will refer to this training phase as Interleaved_S,T1_). To simulate sleep, the rewarded STDP rule was replaced by unsupervised STDP, ensuring a truly offline learning period, and hidden layer neurons were artificially stimulated by Poisson distributed spike trains in order to maintain spiking rates similar to that during task training (indeed, in vivo, activity of the neocortical neurons during REM sleep is similar to awake^32^; see *Methods: Simulated Sleep* for details). Importantly, no training on Task 2 (old task here) was performed at any time during Interleaved_S,T1_. Figure 4B shows that following Interleaved_S,T1_ the network achieved a performance of 0.69 ± 0.02 on Task 1 and a performance of 0.67 ± 0.03 on Task 2, comparable to both single task performances following sequential training on Task 1 (0.70 ± 002) and Task 2 (0.69 ± 0.03) (Figure 1B/C) and exceeding those achieved through Interleaved_T1,T2_ training (Figure 3B). When durations of Task 1 individual training episodes was increased significantly beyond 100 cycles during Interleaved_S,T1_, the network was only able to perform well on the new Task 1 while performance on the old Task 2 dropped to the chance level (not shown).

**Figure 4.**
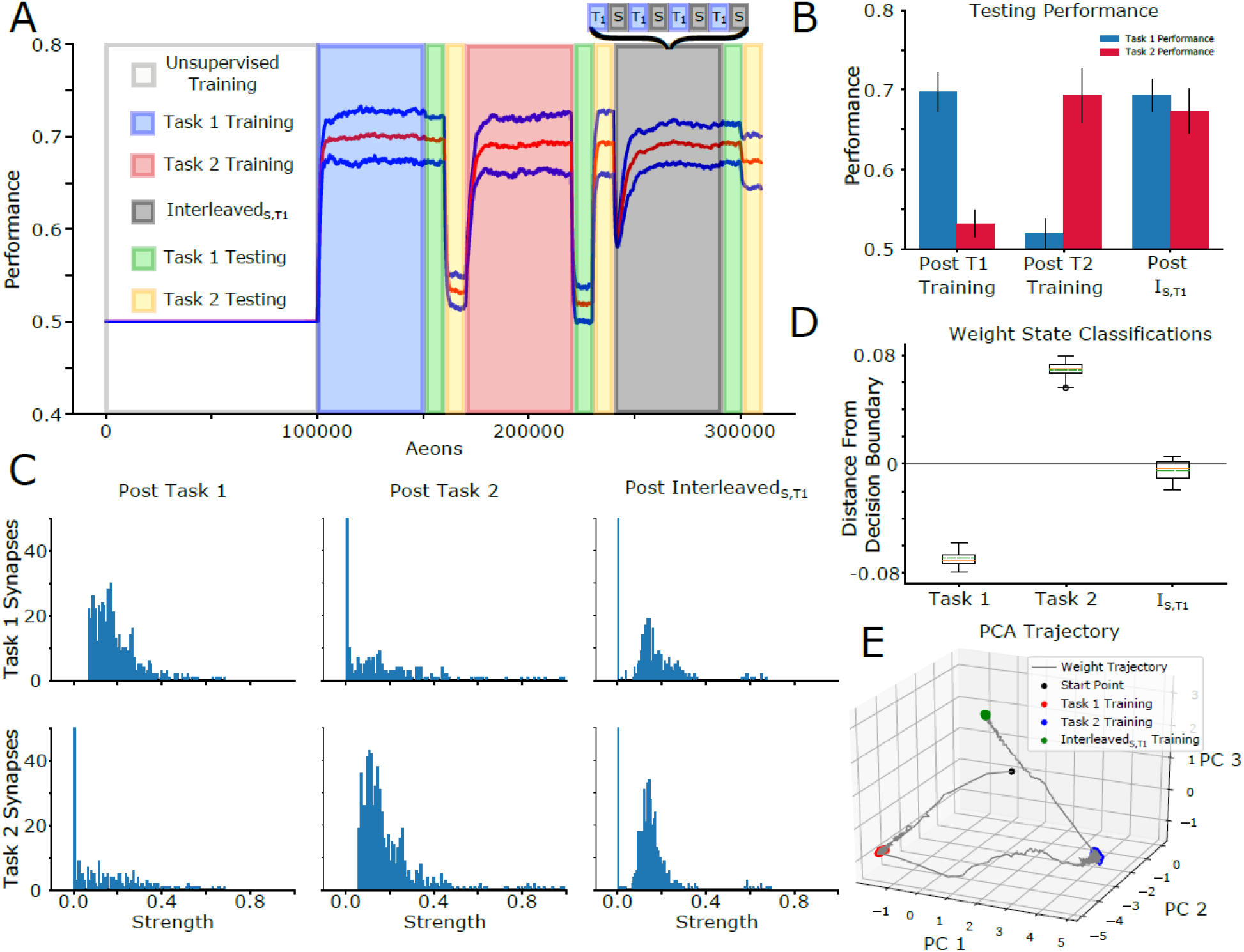
Periods of sleep interleaved with training on a new task can prevent catastrophic forgetting. **(A)** Task paradigm similar to that shown in (3A) but with 50,000 aeons of Interleaved_S,T1_ training (gray) instead of Interleaved_T1,T2_ training. Note that performance for Task 2 remains high despite no Task 2 training during this period. **(B)** Mean and standard deviation of performance during testing on Task 1 (blue) and Task 2 (red) after each training period. Following Task 1 training, mean performance on Task 1 was 0.70 ± 0.02 while Task 2 was 0.53 ± 0.02. Conversely, following Task 2 training, mean performance on Task 1 was 0.52 ± 0.02 while Task 2 was 0.69 ± 0.04. Following Interleaved_S,T1_ training, mean performance on Task 1 was 0.69 ± 0.02 while Task 2 was 0.67 ± 0.03. **(C)** Distributions of task-relevant synaptic weights. The distributional structure of Task 1-relevant synapses following Task 1 training (topleft) is destroyed following Task 2 training (top-middle), but partially recovered following Interleaved_S,T1_ training (top-right). Similarly, the distributional structure of Task 2-relevant synapses following Task 2 training (bottom-middle), which was not present following Task 1 training (bottom-left), was partially preserved following Interleaved_S,T1_ training (bottom-right). Task-relevant synapses were considered to be those which had a synaptic weight of at least 0.1 following training on that task. **(B)** Box plots with mean (dashed green line) and median (dashed orange line) of the distance to the decision boundary found by an SVM trained to classify Task 1 and Task 2 synaptic weight matrices for Task 1, Task 2, and Interleaved_S,T1_ training across trials. Task 1 and Task 2 synaptic weight matrices had mean classification values of −0.069 and 0.069 respectively, while that of Interleaved_S,T1_ training was −0.0047. **(C)** Trajectory of H to O layer synaptic weights through PC space. Synaptic weights which evolved during Interleaved_S,T1_ training (green dots) clustered in a location of PC space intermediary between the clusters of synaptic weights which evolved during training on Task 1 (red dots) and Task 2 (blue dots).

We interpret these results as follows (see sections below for detailed synaptic connectivity analysis). Each episode of new Task 1 training improves Task 1 performance but damages synaptic connectivity responsible for old Task 2. If continuous Task 1 training is long enough, the damage to Task 2 becomes irreversible. Having a sleep phase after a short period of Task 1 training enables spontaneous forward (H->O) replay that preferentially benefits the strongest synapses. Thus, if Task 2 synapses are still strong enough, they are replayed and increase. To keep the protocol consistent with our previous experiments on Interleaved_T1,T2_ training, we used a combination of sleep and Task 1 training – the same task that was initially trained to naïve network but overwritten during Task 2 training (i.e., entire sequence of events was T1 -> T2 -> Interleaved_S,T1_). However, we obtained the same results in an experiment when, after initial Task 1 training, Task 2 training was interleaved with sleep (i.e., T1 -> InterleavedS,T2), which prevented forgetting Task 1 while Task 2 was learned (see Extended Data Figure 1).

We next traced “task-relevant” synapses, i.e. synapses identified in the top 10% distribution following training on that specific task (Figure 4C; compare to Figure 3C for Interleaved_T1,T2_ training). The structure in the distribution of Task 1-relevant synapses following Task 1 training (Figure 4C; top-left) was destroyed following Task 2 training (top-middle) but partially recovered following Interleaved_S,T1_ training (top-right). The structure in the distribution of Task 2-relevant synapses following Task 2 training (bottom-middle) was not present following Task 1 training (bottom-left) and was partially retained following Interleaved_S,T1_ training (bottom-right). Thus, sleep can preserve important synapses while incorporating new ones.

Figure 4D shows that the SVM robustly classified the synaptic weight states from Task 1 and Task 2 while those from Interleaved_S,T1_ weight states fell significantly closer to the decision boundary. This indicates that, similar to Interleaved_T1,T2_, the synaptic weight matrices which result from Interleaved_S,T1_ training are a mixture of Task 1 and Task 2 states. The trajectory of the synaptic weights in PC space shown in Figure 4E provides a visualization of these dynamics. Importantly, the smoothness of this trajectory to its steady state suggests that Task 2 information is never completely erased during this evolution. We take this as evidence that Interleaved_S,T1_ training is capable of integrating synaptic information relevant to Task 1 while preserving Task 2 information.

### Receptive fields of decision-making neurons after sleep represent multiple tasks

To observe that the network has learned both tasks after Interleaved_S,T1_ training, we mapped the receptive fields of decision-making neurons in layer O (Figure 5; see Figure 2 for comparison). Figure 5A shows the receptive field for the neuron in layer O which controls movement in the upper-left direction. This neuron responds to both horizontal (rewarded for Task 1) and vertical (rewarded for Task 2) orientations in the upper-left quadrant of the visual field. Although it initially appears that this layer O neuron may also be responsive to diagonal patterns in this region, analysis of the receptive fields of neurons in layer H (Figure 5B) revealed that these receptive fields are selective to either horizontal food particles (left; rewarded for Task 1) or vertical food particles (right; rewarded for Task 2) in the upper-left quadrant of the visual field. Other receptive fields were responsible for avoidance of punished particles for both tasks (see examples in Figure 5B, bottom-middle-right and bottom-middle-left). Thus, the network will utilize one of two distinct sets of layer H neurons, selective for either Task 1 or Task 2, depending on which food particles are present in the environment.

**Figure 5.**
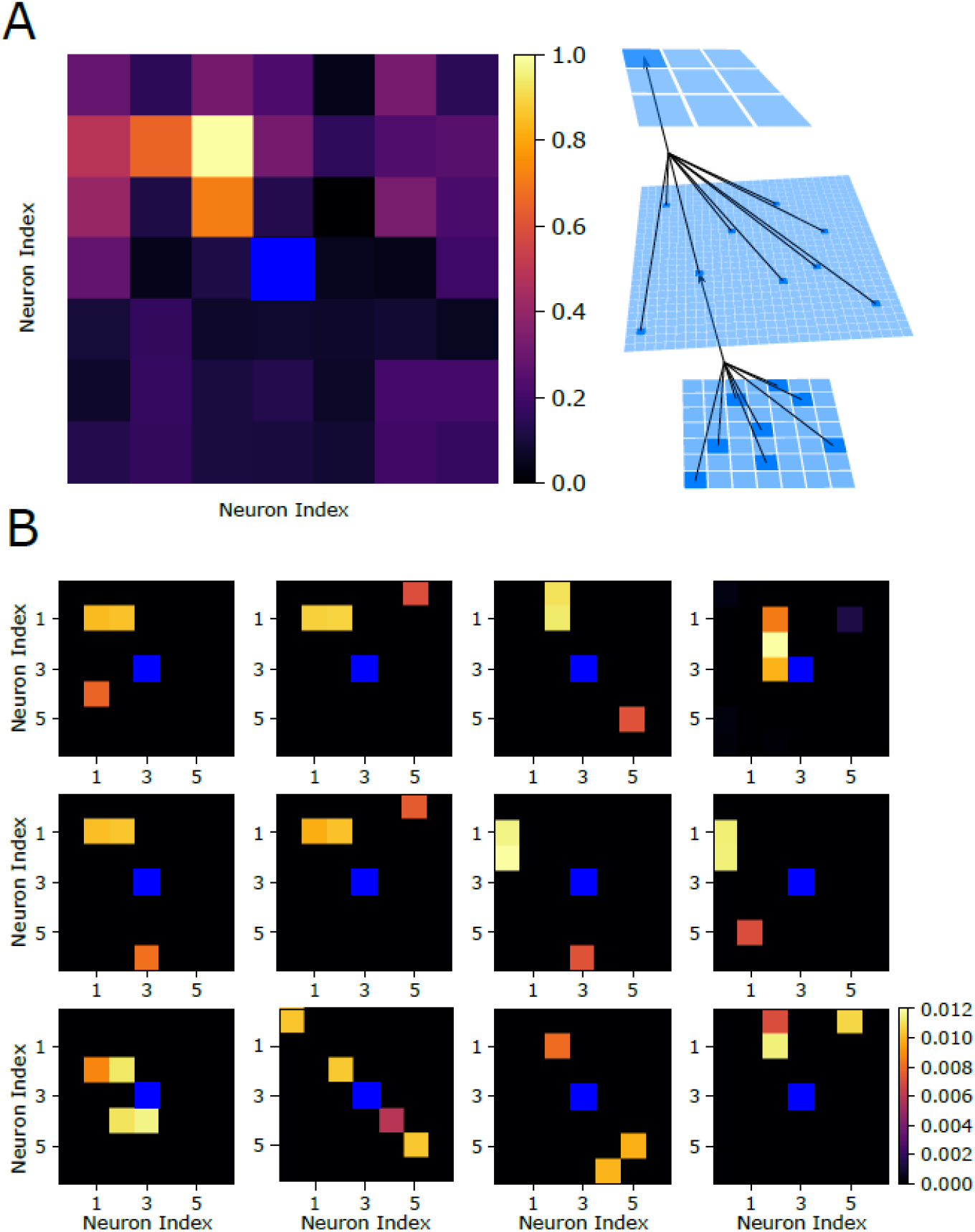
Receptive fields following interleaved Sleep and Task 1 training reveal how the network can multiplex the complementary tasks. **(A)** Left, Receptive field of the output layer neuron controlling movement to the upper-left direction following interleaved sleep and Task 1 training. This neuron has a complex receptive field capable of responding to horizontal and vertical orientations in the upper-left quadrant of the visual field. Right, Schematic of the connectivity between layers. **(B)** Examples of receptive fields of hidden layer neurons which synapse strongly onto the output neuron from (A) after interleaved Sleep and Task 1 training. The majority of these neurons selectively respond to horizontal food particles (left half) or vertical food particles (right half) in the upper-left quadrant of the visual field, promoting movement in that direction and acquisition of the rewarded patters. A few neurons (bottom-middle-left/right) can be seen to selectively respond to the presence of positive/negative diagonal food particles in the bottom-right quadrant of the visual field. Activation of these neurons will promote avoidance movement to the upper-left direction away from the punished patterns.

### Periods of sleep allow reintegration of new task without interference through renormalization of task-relevant synapses

To visualize synaptic weight dynamics during Interleaved_S,T1_ training, traces of all synapses projecting to a single representative output layer neuron were plotted (figure 6A). At the onset of Interleaved_S,T1_ training (i.e. 240,000 aeons), the network was only able to perform on Task 2, meaning the strong synapses in the network were specific to this task. These synapses were represented by a cluster ranging from ~0.08 to ~0.4; the rest of synapses grouped near 0. As Interleaved_S,T1_ training progressed, Task 1 specific synapses moved to the strong cluster and some, presumably less important, Task 2 synapses moved to the weak cluster. After a period of time the rate of transfer decreased and the total number of synapses in each group stabilized, showing that the network is approaching equilibrium (Figure 6B).

**Figure 6.**
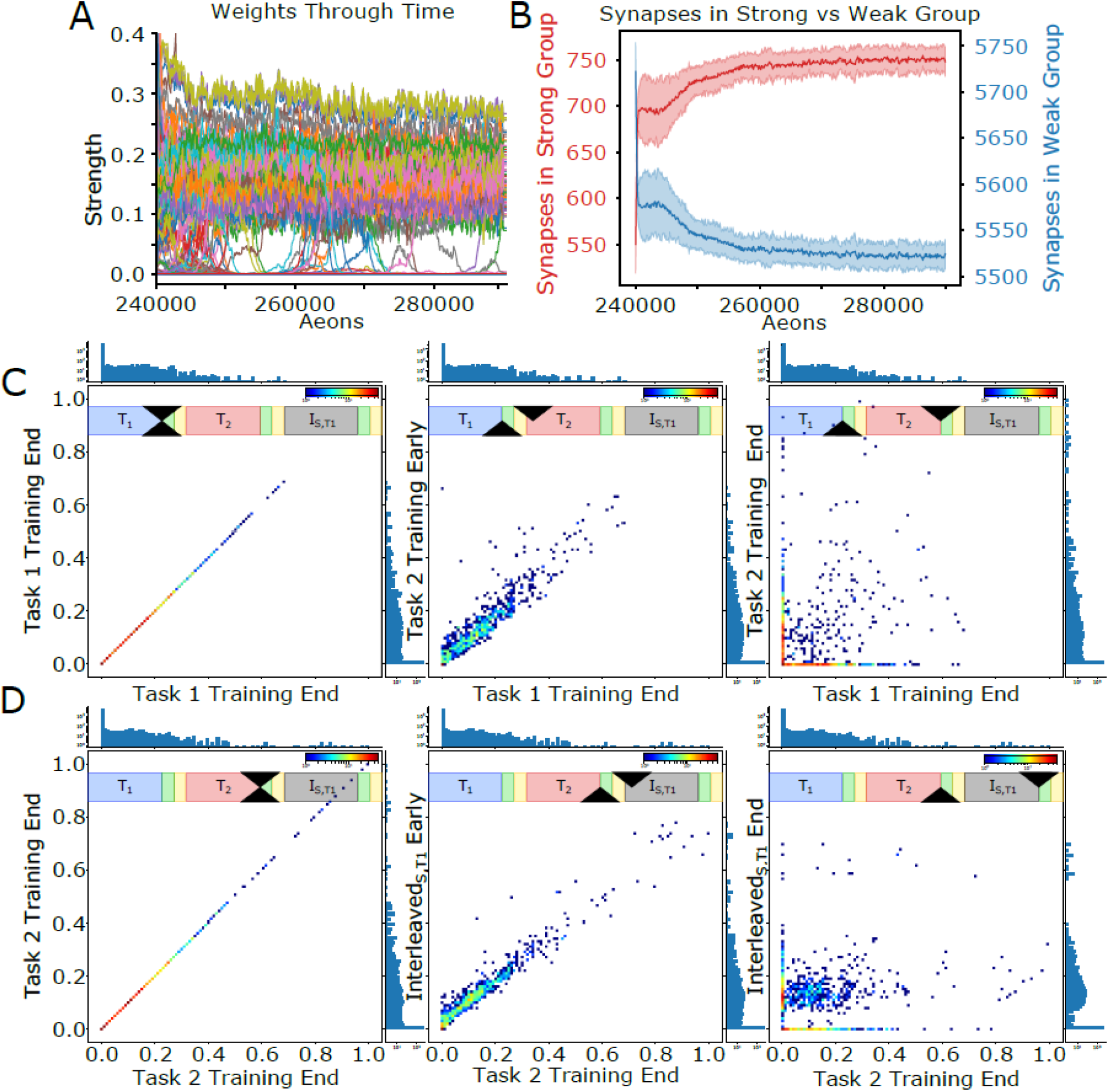
Periods of sleep allow learning Task 1 without interference with old Task 2 through renormalization of task-relevant synapses. **(A)** Dynamics of all incoming synapses to a single output layer neuron during Interleaved_S,T1_ training shows the synapses separate into two clusters. **(B)** Number of synapses in the strong (red) and weak (blue) clusters during Interleaved_S,T1_. **(C)** Two-dimensional histograms illustrating synaptic weights dynamics. For each plot, the x-axis represents synaptic weight after Task 1 training and the y-axis represents the synaptic weight at a different point in time(Scale bar: brown - 50 synapses/bin, blue - 1 synapse/bin. One-dimensional projections along top and right sides show the global distribution of synapses at the time slices for a given plot. If no training occurred between the time slices, a diagonal plot depicts that synaptic weights have not changed (left). After a small amount of Task 2 training, all points lie near the diagonal (middle) indicating minimal changes to synaptic weights. Once Task 2 is fully trained (right), many synapses move far away from their original values. In particular, a red cluster along the x-axis indicates synapses which were strong after Task 1 training but were erased after Task 2 training. **(D)** Same as (C) except the x-axis refers to the end of Task 2 training. Again, a diagonal plot is attained when no training takes place between the time slices (left), and points lie near the diagonal when only a small amount of Interleaved_S,T1_ training occurs (middle). After a full period of Interleaved_S,T1_ training (right), weak synapses were recruited to support Task 1 (red cluster along the y-axis) and many Task 2 specific synapses remained moderately strong (blue cluster along x-axis).

To visualize how sleep renormalizes task relevant synapses, we plotted two-dimensional weight distributions for Task 1->Task2 (Figure 6C) and Task 2 -> Interleaved_S,T1_ (Figure 6D) experiments (see *Methods: 2-D Synaptic Weight Distributions* for details). To establish a baseline, in Figure 6C (left) the weight state at the end of Task 1 training (X-axis) (see overall timeline of this experiment in Figure 4A) was compared to itself (Y-axis). This formed a perfectly diagonal plot. Most synapses were weak (red dots) with stronger synapses forming a tail in the distribution. The next comparison (Figure 6C, middle) was between the weight state after Task 1 training (X-axis) and a time early on Task 2 training (Y-axis). At that time, synapses were only able to modify their strength slightly, causing most points to lie close to the diagonal. As training on Task 2 continued until maximum performance was reached, synapses moved far away from the diagonal (Figure 6C, right). Two trends were observed. A set of synapses that had a strength near zero following Task 1 training increased strength following Task 2 training (Figure 6D, right, red dots along Y-axis). At the same time, many strongly trained by Task 1 synapses were depressed down to zero (Figure 6C, right, red dots along X-axis). The latter illustrates the effect of catastrophic forgetting - complete overwriting of the synaptic weight matrix caused performance of Task 1 to return to baseline after training on Task 2.

Does sleep prevent overwriting of the synaptic weight matrix? The Figure 6D plots use the weight state at the end of training Task 2 as a reference that is compared to different times during Interleaved_S,T1_ training. The first two plots (Figure 6D, left/middle) are similar to those in Figure 6C. However, after Interleaved_S,T1_ training (Figure 6D, right) many synapses that were strong following Task 2 training were not depressed to zero but rather were pushed to an intermediate strength where they are still functional (note cluster of points parallel to X-axis; see also projection to 1D on the right side of the graph). Thus, Interleaved_S,T1_ training, combining new training on Task 1 with periods of unsupervised sleep, moved synapses in a way that preserved strong synapses from a previously learned task while also introducing new strong synapses to perform a new task. Since a significant fraction of the strong synapses from training on Task 2 were preserved (due to the sleep periods), performance on Task 2 remained high following Interleaved_S,T1_ training despite the fact that the networks received no new training examples of Task 2.

### Periods of sleep push the network towards the intersection of the solution manifolds representing Task 1 and Task 2 specific weight configurations

To visualize the approximate task-specific solution manifolds (*M_T1_* and *M_T2_*) and their intersection (*M_T1∩T2_*) in synaptic weight space, we used multiple trials (with different initialization) of Task 1 and Task 2 training to sample the manifolds. Figure 7A shows (in kPCA space) that multiple different configurations of synaptic weights can provide high performance for a given task. For example, all red dots in Figure 7A represent the states with the same high level of performance for Task 1 (but not Task 2). In addition, cyan and green dots represent states with high level of performance for both Task 1 and Task2. We interpret these results as evidence that synaptic weight space includes a manifold, *M_T1_*, where different configurations of weights (red, green, cyan dots) all allow for Task 1 to perform well. This manifold intersects with another one, *M_T2_*, where different weights configurations (blue, green, cyan dots) are all suitable for Task 2. Figures 7B and 7C show 2D projections of this space onto PCs 1 and 2 and PCs 1 and 3, respectively. From these projections, we can see that PC 1 seems to capture the extent to which a synaptic weight configuration is associated with Task 1 (positive values) or Task 2 (negative values), while PC 2 and PC 3 capture the variance in synaptic weight configurations associated with Task 1 and Task 2, respectively. Note, the trajectories through this space (red/blue lines) during Interleaved_T1,T2_ and Interleaved_S,T1/T2_ training would also belong to the respective task manifolds as performance on the old tasks was never lost in these training scenarios.

**Figure 7.**
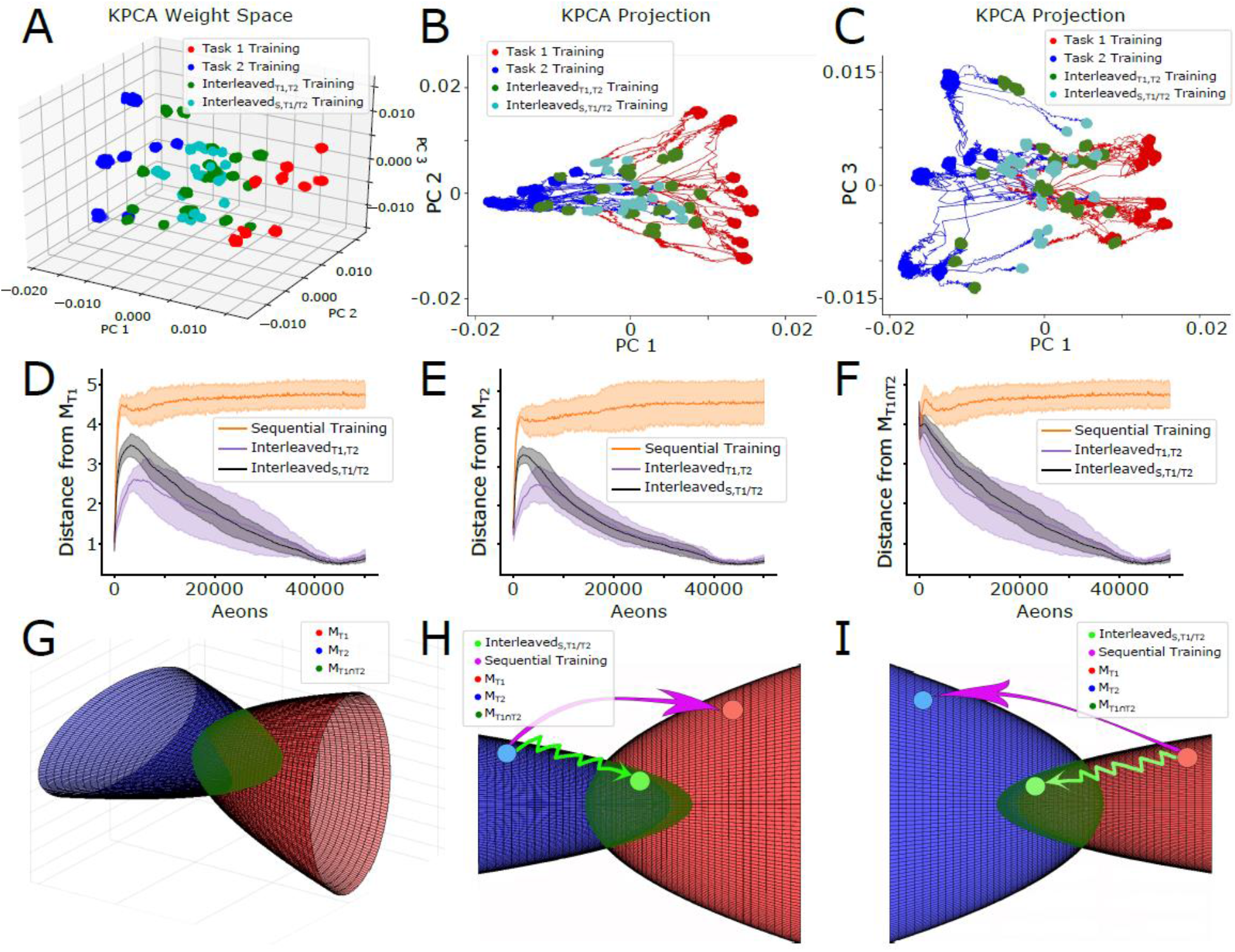
Periods of sleep push the network towards the intersection of Task 1 and Task 2 specific solution manifolds. **(A-C)** Low-dimensional visualizations of the synaptic weight configurations of 10 networks obtained through kPCA for 3-dimensions (A), 2-dimensions using PC 1 and PC 3 (B), and 2-dimensions using PC 1 and PC 3 (C). Synaptic weight configurations taken from the last fifth of Task 1 (red dots), Task 2 (blue dots), Interleaved_T1,T2_ (green dots), and Interleaved_S,T1/T2_ (cyan dots) training are shown. PC 1 characterizes good performance on Task 1 (positively valued) or Task 2 (negatively valued) training. PC 2 (PC 3) characterizes the variability in Task 1 (Task 2) training. Trajectories resulting from Interleaved_T1,T2_ and Interleaved_S,T1/T2_ training following Task 1 (Task 2) training are shown in red (blue). **(D-F)** Average (solid lines) and standard deviation (shaded regions) of the n-dimensional Euclidean distances between the current synaptic weight configuration and *M_T1_* (D), *M_T2_* (E), and *M_T1∩T2_* (F) during Sequential (orange), Interleaved_T1,T2_ (purple), and Interleaved_S,T1/T2_ (black) training. Following Task 2 (D) or Task 1 (E) training, Sequential training on the opposite task causes the synaptic weight configuration to diverge from the initial solution manifold, while Interleaved_T1,T2_ and Interleaved_S,T1/T2_ training do not. (F) Interleaved_T1,T2_ and Interleaved_S,T1/T2_ training cause the synaptic weight configuration to converge to *M_T1∩T2_* while Sequential training avoids this intersection. **(G)** Authors’ interpretation of the task-specific point-sets shown in (A-C) as solution manifolds *M_T1_* (red) and *M_T2_* (blue). *M_T1_* and *M_T2_* can be thought of as two oppositely oriented elliptic paraboloids which intersect orthogonally near the origin *(M_T1∩T2_*; dark green). **(H,I)** Sequential training (pink arrow) causes the network to jump from one solution manifold to the other while avoiding *M_T1∩T2_*, while Interleaved_S,T1/T2_ training (light green arrow) keep the network close to the initial solution manifold as it converges towards *M_T1∩T2_*.

We calculated the distance from the current synaptic weight configurations to *M_T1_* (Figure 7D), *M_T2_* (Figure 7E), and *M_T1∩T2_* (Figure 7F; see *Methods: Distance from Solution Manifolds* for details). Figures 7D and 7E show that while Sequential (T1->T2 or T2->T1) training causes synaptic weight configurations to diverge quickly from its initial solution manifold (i.e. *M_T1_* or *M_T2_*), both Interleaved_T1,T2_ and Interleaved_S,T1/T2_ training cause synaptic weight configurations to stay close to the initial solution manifold as the new task was learned. (Note, that we under sampled *M_T1_* and *M_T2_*, which explains initial distance increase.) Importantly, Figure 7F shows that while both Interleaved_T1,T2_ and Interleaved_S,T1/T2_ training cause synaptic weight configurations to smoothly converge towards *M_T1⋂T2_*, Sequential training avoids this intersection entirely.

In Figure 7G we show a schematic depiction of these results. The task-specific manifolds, *M_T1_* and *M_T2_*, are roughly defined in 3D projection by two orthogonal elliptic paraboloids with opposite orientation, with an approximately ellipsoidal intersection, *M_T1∩T2_*. Figures 7H and 7I depict the trajectories the network takes in this space following Task 2 and Task 1 training, respectively. Sequential training causes the network to jump directly from one task-specific solution manifold to the other, resulting in catastrophic forgetting. In contrast, interleaving new task training with sleep (Interleaved_S,T1/T2_) prevents catastrophic forgetting by keeping the network close to the old task solution manifold as it converges towards *M_T1⋂T2_* – a region capable a supporting both tasks simultaneously.

## Discussion

In this study we report that a multi-layer SNN utilizing reinforcement learning may exhibit catastrophic forgetting upon sequential training of two complementary complex foraging tasks, but the problem is mitigated if the network is allowed, during new task training, to undergo intervening periods of spontaneous reactivation which we consider to be equivalent to the replay observed during periods of sleep in biological systems. This scenario was effectively equivalent to explicit interleaved training of both tasks, however, no training data for the old task were required during “sleep”. At the synaptic level, training a new task alone led to complete overwriting of synaptic weights responsible for the previous task. In contrast, interleaving periods of reinforcement learning on a new task with periods of unsupervised learning during sleep preserved old task synapses damaged by new task training to avoid forgetting and enhanced new task synapses to allow new task learning. Thus, the network was pushed towards the intersection of the solution manifolds representing synaptic weight configurations associated with each task - an optimal compromise for performing both tasks.

The critical role that sleep plays in learning and memory is supported by a vast, interdisciplinary literature spanning both psychology and neuroscience^21,33–36^. Specifically, it has been suggested that REM sleep supports the consolidation of non-declarative or procedural memories while non-REM sleep supports the consolidation of declarative memories^21,35,37^. In particular, REM sleep has been shown to be important for the consolidation of memories of tasks involving perceptual pattern separation, such as the texture discrimination task^21,38^. Despite the difference in the cellular and network dynamics during these two stages of sleep^21,35^, both are thought to contribute to memory consolidation through repeated reactivation, or replay, of specific memory traces acquired during learning^19–21,33,34,37,39^. These studies suggest that through replay, sleep can support the process of off-line memory consolidation to circumvent the problem of catastrophic forgetting.

From mechanistic perspective, the sleep phase in our model protects old memories by enabling unsupervised learning - spontaneous replay of synapses responsible for previously learned tasks. We previously reported that in the thalamocortical models, a sleep phase may enable replay of spike sequences learned in awake to improve post-sleep performance^39,40^ and to protect old memories from catastrophic forgetting^41^. Although in this work we model sleep and noise with spiking statistics similar to awake training, theoretical work from another group has also shown that noise causes implicit rehearsal of older memories which protects against interference^42^. Here we found, however, that a single episode of new task training using reinforcement learning could quickly erase an old memory to the point that it cannot be recovered by subsequent sleep. The solution was similar to how brain slowly learns procedural (hippocampal-independent) memories^21,35,37,38,43^. Each episode of new task training improves this task performance only slightly but also damages slightly synaptic connectivity responsible for the older task. Subsequent sleep phases enable replay that preferentially benefits the strongest synapses, such as those from old memory traces, to allow them to recover.

We found that multiple distinct configurations of synaptic weights can support each task in our model, suggesting the existence of task specific solution manifolds in synaptic weight space. Sequential training of new tasks makes the network to jump from one solution manifold to another, enabling memory for the most recent task but erasing memories of the previous tasks. Interleaving new task training with sleep phases enables the system to evolve towards intersection of these manifolds where synaptic weight configurations can support multiple tasks (a similar idea was recently proposed in the machine learning literature to minimize catastrophic interference by learning representations that accelerate future learning^44^). From this point of view having multiple episodes of new task training interleaved with multiple sleep episodes allows gradual convergence to the intersection of the manifolds representing old and new tasks, while a single long episode of new task learning would push the network far away from the old task manifold making it impossible to recover by subsequent sleep.

Although classical interleaved training showed similar performance results in our model as interleaving training with sleep, we believe the latter to be superior on the following theoretical grounds. Classical interleaved training will necessarily cause the system to oscillate about the optimal location in synaptic weight space which can support both tasks because each training cycle uses a cost function specific to only a single task. While this can be ameliorated with a learning rate decay schedule, the system is never actually optimizing for the desired dualtask state. Sleep, on the other hand, can support not only replays of the old task, but also support replays which are a mixture of both tasks^42,45,46^. Thus, through unsupervised learning during sleep replay, the system is able to perform approximate optimization for the desired dual-task state.

While our model represents a dramatic simplification of any biological system, we believe that it captures some important processing steps of how animal and human brains interact with the external world. The primary visual system is believed to employ a sequence of processing steps when visual information is increasingly represented by neurons encoding higher level features^28–30^. This processing step was reduced to very simple convolution from input to hidden layer in our model. Subsequently, in the brain, associative areas and motor cortex are trained to make decisions based on reward signals released by neuromodulatory centers^8,47–49^. This was reduced in our model to synaptic projections from the hidden to output (decision making) layer implementing rewarded STDP to learn a task^24–26^.

Our results are in line with a large body of literature suggesting that interleaved training is capable of mitigating catastrophic forgetting in ANNs^4,8,9^ and SNNs^10,11^. The novel contribution from this study is that the data intensive process of interleaved training can be avoided in SNNs by inserting periods of noise-induced spontaneous reactivation – unsupervised learning – during new task training; similar to how brains undergo offline consolidation periods during sleep resulting in reduced retroactive interference to previously learned tasks^21,43^. In fact, our results are in line with previous work done in humans showing that perceptual learning tasks are subject to retroactive interference by competing memories without an intervening period of REM sleep^37,38^. Moreover, performance on visual discrimination tasks in particular have been shown to steadily improve over successive nights of sleep^38^, consistent with our findings that interleaving multiple periods of sleep with novel task learning leads to optimal performance on each task.

Our study predicts synaptic level mechanisms of how sleep-based memory reactivation can protect old memory traces during training of a new interfering memory task. It suggests the apparent loss of recall performance for older tasks in ANNs and SNNs after new training does not necessarily imply a complete erasure of the old task, but instead indicates that the old tasks decision states became unreachable by the associated inputs. Sleep can reverse the damage to synaptic connectivity by replaying the old memory traces without explicit usage of the old training data.

## Methods

### Environment

Foraging behavior took place in a virtual environment consisting of a 50×50 grid with randomly distributed “food” particles. Each particle was two pixels in length and could be classified into one of four types depending on its orientation: vertical, horizontal, positively sloped diagonal, or negatively sloped diagonal. During the initial unsupervised training period, the particles are distributed at random with the constraints that each of the four types are equally represented and no two particles can be directly adjacent. During training and testing periods only the task-relevant particles were present. When a particle was acquired as a result of the virtual agent moving, it was removed from its current location (simulating consumption) and randomly assigned to a new location on the grid, again with the constraint that it not be directly adjacent to another particle. This ensures a continuously changing environment with a constant particle density. The density of particles in the environment was set to 10%. The virtual agent can see a 7×7 grid of squares (the “visual field”) centered on its current location and it could move to any adjacent square, including diagonally, for a total of eight directions.

### Network structure

The network was composed of 842 spiking map-based neurons (see *Methods: Map-based neuron model* below)^50,51^, arranged into three feed-forward layers to mimic a basic biological circuit: a 7×7 input layer (I), a 28×28 hidden layer (H), and a 3×3 output layer (O) with a nonfunctional center neuron (Fig 1). Input to the network was simulated as a set of suprathreshold inputs to the neurons in layer I, equivalent to the lower levels of the visual system, which represent the position of particles in an egocentric reference frame relative to the virtual agent (positioned in the center of the 7×7 visual field). The most active neuron in layer O, playing the role of biological motor cortex, determined the direction of the subsequent movement. Each neuron in layer H, which can be loosely defined as higher levels of the visual system or associative cortex, received excitatory synapses from 9 randomly selected neurons in layer I. These connections initially had random strengths drawn from a normal distribution. Each neuron in layer H connected to every neuron in layer O with both an excitatory (*W_ij_*) and an inhibitory (*WI_ij_*) synapse. This provided an all-to-all connectivity pattern between these two layers and accomplished a balanced feed-forward inhibition^52^ found in many biological structures^52–57^. Initially, all these connections had uniform strengths and the responses in layer O were due to the random synaptic variability. Random variability was a property of all synaptic interactions between neurons and was implemented as variability in the magnitude of the individual synaptic events.

### Policy

Simulation time was divided up into epochs of 600 timesteps, each roughly equivalent to 300 ms. At the start of each epoch the virtual agent received input corresponding to locations of nearby particles within the 7×7 “visual field”. Thus 48 of the 49 neurons in layer I received input from a unique location relative to the virtual agent. At the end of the epoch the virtual agent made a single move based on the activity in layer O. If the virtual agent moved to a grid location with a “food” particle present, the particle was removed and assigned to a randomly selected new location.

Each epoch was of sufficient duration for the network to receive inputs, propagate activity forward, produce outputs, and return to a resting state. Neurons in layer I which represent locations in the visual field containing particles received a brief pulse of excitatory stimulation sufficient to trigger a spike; this stimulation was applied at the start of each movement cycle (epoch). At the end of each epoch the virtual agent moved according to the activity which has occurred in layer O.

The activity in layer O controlled the direction of the virtual agent’s movement. Each of the neurons in layer O mapped onto a specific direction (i.e. one of the eight adjacent locations or the current location). The neuron in layer O which spiked the greatest number of times during the first half of the epoch defined the direction of movement for that epoch. If there was a tie, the direction was chosen at random from the set of tied directions. If no neurons in layer O spiked, the virtual agent continued in the direction it had moved during the previous epoch.

There was a 1% chance on every move that the virtual agent would ignore the activity in layer O and instead move in a random direction. Moreover, for every movement cycle that passed without the virtual agent acquiring a particle, this probability was increased by 1%. The random variability promoted exploration vs exploitation dynamics and essentially prevented the virtual agent from getting stuck in movement patterns corresponding to infinite loops. While biological systems could utilize various different mechanisms to achieve the same goal, the method we implemented was efficient and effective for the scope of our study.

### Neuron models

For all neurons we used spiking model identical to the model used in^12,13^ that can be described by the following set of difference equations^51,58,59^:

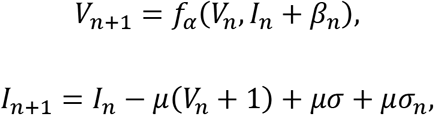

where *V_n_* is the membrane potential, *I_n_* is a slow dynamical variable describing the effects of slow conductances, and *n* is a discrete time-step (0.5 ms). Slow temporal evolution of *I_n_* was achieved by using small values of the parameter *μ* << 1. Input variables *β_n_* and *σ_n_* were used to incorporate external current 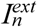 (e.g. background synaptic input): 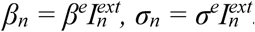. Parameter values were set to *σ* = 0.06, *β^e^* = 0.133, *σ^e^* = 1, and *μ* = 0.0005. The nonlinearity *f_α_*(*V_n_, I_n_*) was defined in the form of the piece-wise continuous function:

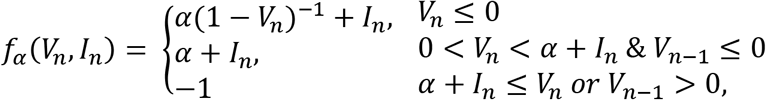

where *α* = 3.65.

This model is very computationally efficient, and, despite its intrinsic low dimensionality, produces a rich repertoire of dynamics capable of mimicking the dynamics of Hodgkin-Huxley type neurons both at the single neuron level and in the context of network dynamics^51,58,60^.

To model the synaptic interactions, we used the following piece-wise difference equation:

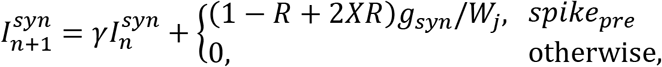

Here *g_syn_* is the strength of the synaptic coupling, modulated by the target rate *W_j_* of receiving neuron *j*. Indices *pre* and *post* stand for the pre- and post-synaptic variables, respectively. The first condition, *spike_pre_*, is satisfied when the pre-synaptic spikes are generated. Parameter *γ* controls the relaxation rate of synaptic current after a presynaptic spike is received (0 ≤ *γ* < 1). The parameter *R* is the coefficient of variability in synaptic release. The standard value of *R* is 0.12. *X* is a random variable sampled from a uniform distribution with range [-1, 1]. Parameter *V_rp_* defines the reversal potential and, therefore, the type of synapse (i.e. excitatory or inhibitory). The term (1-*R*+2*XR*) introduces a variability in synaptic release such that the effect of any synaptic interaction has an amplitude that is pulled from a uniform distribution with range [1-*R*, 1+*R*] multiplied by the average value of the synapse.

### Synaptic plasticity

Synaptic plasticity closely followed the rules introduced in^12,13^. A rewarded STDP rule^24–27^ was operated on synapses between layers H and O while a standard STDP rule operated on synapses between layers I and H. A spike in a post-synaptic neuron that directly followed a spike in a pre-synaptic neuron created a *pre before post* event while the converse created a *post before pre* event. Each new post-synaptic (pre-synaptic) spike was compared to all pre-synaptic (post-synaptic) spikes with a time window of 120 iterations.

The value of an STDP event (trace) was calculated using the following equation^22,23^:

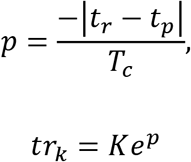

where *t_r_* and *t_p_* are the times at which the pre- and post-synaptic spike events occurred respectively, *T_c_* is the time constant and is set to 40 ms, and *K* is maximum value of the trace *tr_k_* and is set to −0.04 for a *post before pre* event and 0.04 for a *pre before post* event.

A trace was immediately applied to synapse between neurons in layers I and H. However, for synapses between neurons in layers H and O the traces were stored for 6 epochs after its creation before being erased. During storage, a trace had an effect whenever there was a rewarding or punishing event. In such a case, the synaptic weights are updated as follows:

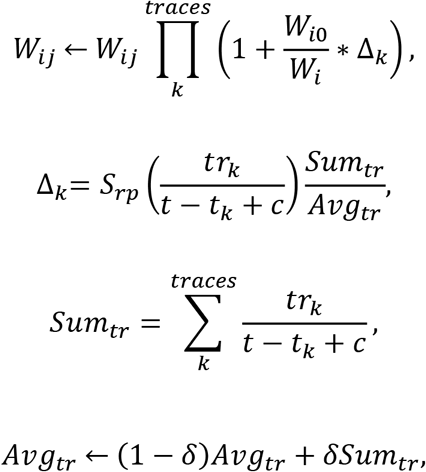

where *t* is the current timestep, *S_rp_* is a scaling factor for reward/punishment, *tr_k_* is the magnitude of the trace, *t_k_* is the time of the trace event, *c* is a constant (=1 epoch) used for decreasing sensitivity to very recent spikes, *W_i_* = *Σ_j_ W_ij_* is the total synaptic strength of all connections from the neuron *i* in layer H to all neurons in layer O, *W_i0_* is a constant that is set to the initial value *(target value)* of *W_i_* at the beginning of the simulation. The term *W_i0_/W_i_* helped to keep the output weight sum close to the initial target value. The effect of these rules was that neurons with lower total output strength could increase their output strength more easily.

The network was rewarded when the virtual agent moved to a location which contained a particle from a “food” pattern (horizontal in Task 1, vertical in Task 2) and *S_rp_* = 1, and received a punishment of *S_rp_* = −0.001 when it moved to a location with a particle from a neutral pattern (negative/positive diagonal in Task 1/2). A small punishment of *S_rp_* = −0.0001 was applied if the agent moved to a location without a particle present to help the virtual agent learn to acquire “food” as rapidly as possible. During periods of sleep the network received a constant reward of *S_rp_* = 0.5 on each movement cycle.

To ensure that neurons in layer O maintained a relatively constant long-term firing rate, the model incorporated homeostatic synaptic scaling which was applied every epoch. Each timestep, the total strength of synaptic inputs *W_j_* = *Σ_i_ W_ij_* to a given neuron in layer O was set equal to the target synaptic input *W_j0_* – a slow variable which varied over many epochs depending on the activity of the given neuron in layer O – which was updated according to:

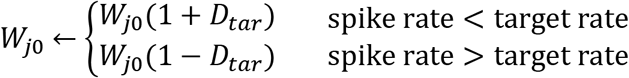

To ensure that the net synaptic input *W_j_* to any neuron was unaffected by plasticity events at the individual synapses at distinct timesteps and equal to *W_j0_*, we implemented a scaling process akin to heterosynaptic plasticity which occurs after each STDP event. When any excitatory synapse of neuron in layer O changed in strength, all other excitatory synapses received by that neuron were updated according to:

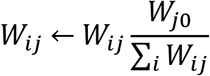

### Simulated Sleep

To simulate the sleep phase, we inactive the sensory receptors (i.e. the input layer of network), cut off all sensory signals (i.e. remove all particles from the environment), and decouple output layer activity from motor control (i.e. the output layer can spike but no longer causes the agent to move). We also change the learning rule between the hidden and output layer from rewarded to unsupervised STDP (see *Methods: Synaptic Plasticity* for details) as there is no way to evaluate decision-making without sensory input or motor output.

To simulate the spontaneous activity observed during REM sleep, we provided noise to each neuron in the hidden layer in a way which ensured that the spiking statistics of each neuron was conserved across awake and sleep phases. To determine these spiking rates, we recorded average spiking rates of neurons in the hidden layer H during preceding training of both Task 1 and Task 2; these task specific spiking rates were then averaged to generate target spiking rates for hidden layer neurons. Interleaved_S,T1_ training consisted of alternating intervals of this sleep phase and training on Task 1, with each interval lasting 100 movement cycles (although no movement occurred).

### Support Vector Machine Training

A support vector machine with a radial basis function kernel was trained to classify synaptic weight configurations as being related to Task 1 or Task 2. Labeled training data were obtained by taking the excitatory synaptic weight matrices between the hidden and output layers from the last fifth of the Task 1 and Task 2 training phases (i.e. after performance had appeared to asymptote). These synaptic weight matrices were then flattened into column vectors, and the column vectors were concatenated to form a training data matrix of size *number of features* x *number of samples.* The number of features was equal to the total number of excitatory synapses between the hidden and output layer – 6272 dimensions. We then used this support vector machine to classify held out synaptic weight configurations from Task 1 and Task 2 training, as well as ones which resulted from Interleaved_T1,T2_ and Interleaved_S,T1_ training.

### 2-D Synaptic Weight distributions (Figure 6)

First for each synapse we found how its synaptic strength changes between two slices in time, where the given synapse’s strength at time slice 1 is the point’s X-value and strength at time slice 2 is its Y-value. Then we binned this space and counted synapses in each bin to make two dimensional histograms where blue color corresponds to a single synapse found in a bin and brown corresponds to the max of 50 synapses. These two-dimensional histograms assist in visualizing the movement of all synapses between the two slices in time that are specified by the timelines at the top of each plot. Conceptually, it is important to note that if a synapse does not change in strength between time slice 1 and time slice 2, then point the synapse corresponds to in this space will lie on the diagonal of the plot since the X-value will match the Y-value. If a great change in the synapse’s strength has occurred between time slice 1 and time slice 2, then the synapse’s corresponding point will lie far from the diagonal since the X-value will be distant from the Y-value. The points on the X-(Y-) axis represent synapses that lost (gained) all synaptic strength between time slice 1 and time slice 2.

### Distance from Solution Manifolds (Figure 7)

Each of the two solution manifolds (i.e. Task 1 and Task 2 specific manifolds) were defined by the point-sets in synaptic weight space which were capable of supporting robust performance on that particular task, namely the sets *M_T1_* and *M_T2_*. This included the synaptic weight states from the last fifth of training on a particular task (i.e. after performance on that task appeared to asymptote) and all of the synaptic weight states from the last fifth of both Interleaved_T1,T2_ and Interleaved_S,T1/T2_ training. The intersection of the two solution manifolds (i.e. the point-set *M_T1∩T2_*) was defined solely by the synaptic weight states from the last fifth of both Interleaved_T1,T2_ and Interleaved_S,T1_ training. As the network evolved along its trajectory in synaptic weight space, the distance from the current point in synaptic weight space, *p_t_*, to the two solution manifolds and their intersection were computed as follows:

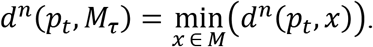

Here, *d^n^* is the n-dimensional Euclidean-distance function, where *n* is the dimensionality of synaptic weight space (i.e. *n* = 6272 here), *M_τ_* is the point-set specific to the manifold or intersection in question (i.e. either *M_T1_, M_T2_*, or *M_T1⋉T2_*), and *x* is a particular element of the point-set *M*.

## Acknowledgements

This study was supported by Lifelong Learning Machines program from DARPA/MTO (HR0011-18-2-0021) and ONR MURI (N000141612829)

## Author Contributions

J.E.D. carried out experiments; J.E.D conducted computational analyses; R.G. designed experiments; R.G. and J.E.D. designed analysis approaches; R.G., J.E.D., P.S., and M.B. interpreted data; R.G., P.S., and M.B. conceived the project; R.G. and M.B. wrote the majority of the paper. All authors commented on and contributed to the paper.

---

**Extended Data Figure 1.**
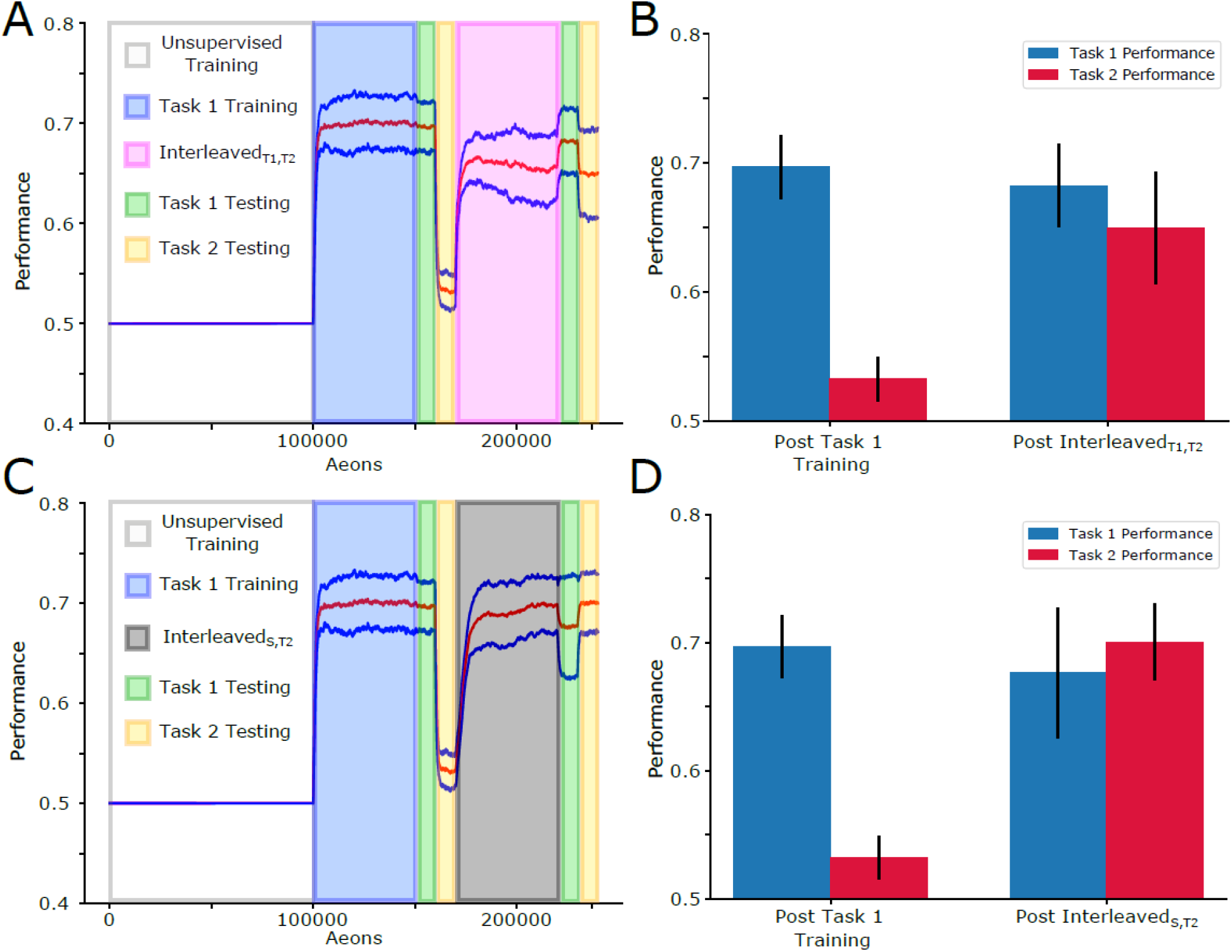
Interleaved training of two tasks and interleaving training new task with sleep both can integrate new tasks without catastrophic forgetting. **(A).** Mean performance (red line) and standard deviation (blue lines) over time: 100,000 aeons of unsupervised training (white), 50,000 aeons of Task 1 training (blue), 10,000 aeons of Task 1 (green) and Task 2 (yellow) testing, 50,000 aeons of Interleaved_T1,T2_ training (pink), 10,000 aeons of Task 1 (green) and Task 2 (yellow) testing. **(B)** Mean and standard deviation of performance during testing on Task 1 (blue) and Task 2 (red) after each training period. Following Task 1 training, mean performance on Task 1 was 0.69 ± 0.02 while Task 2 was 0.53 ± 0.02. Following Interleaved_T1,T2_ training, mean performance on Task 1 was 0.68 ± 0.03 while Task 2 was 0.64 ± 0.04. **(C)** Task paradigm similar to that shown in (A) but with 50,000 aeons of InterleavedS,T2 training (gray) instead of Interleaved_T1,T2_ training. **(D)** Mean and standard deviation of performance during testing on Task 1 (blue) and Task 2 (red) after each training period. Following Task 1 training, mean performance on Task 1 was 0.70 ± 0.02 while Task 2 was 0.53 ± 0.02. Following Interleaved_S,T2_ training, mean performance on Task 1 was 0.68 ± 0.05 while Task 2 was 0.70 ± 0.03.

